# Sexually antagonistic co-evolution at the molecular level: patterns of genetic co-variation predict phenotypic outcomes of mating interactions

**DOI:** 10.1101/2025.11.04.686516

**Authors:** Alma Thorson, Rhonda R. Snook, R. Axel W. Wiberg

## Abstract

Sexually antagonistic co-evolution (SAC) is a potent driver of phenotypic change, with males and females engaged in cycles of co-evolution in which sex-specific trait expression enhances their own fitness at the expense of the other. The cyclical process results in transient mismatches between male and female traits wherein the relative exaggeration of sex-specific phenotypes predicts which sex has the upper-hand during mating interactions. SAC should also result in genetic signatures of co-evolution, and relative mismatches in differentiation between the genetic components of male- and female- conflict traits should predict outcomes of mating interactions. However, due to a poor knowledge of genes underlying SAC traits these predictions remain untested, impeding understanding of how SAC shapes genetic variation and the phenotypic consequences. Here we show the first evidence that SAC theory readily extends to the molecular level and classic patterns of phenotypic SAC are recapitulated. We exploit the iconic sex-peptide (SP) network in *Drosophila melanogaster*, which comprises male seminal fluid proteins and female reproductive tract proteins that interact to influence female post-mating behaviours. Using genomic data from ∼150 populations, we show that genetic differentiation at loci of the male- and female- components of the SP-network loci co-varies as expected under SAC. We use relative levels of genetic differentiation between male- and female- components of the SP-network to infer which populations should have male- or female- advantage in mating interactions. Mating and survival assays using replicated lab populations established from wild-derived isofemale lines show populations with inferred male-advantage have reduced female remating and earlier peaks of egg production compared to populations with inferred female-advantage. Thus, relative levels of genetic differentiation in the classic SP-network with known SAC phenotypes accurately predicts outcomes of mating interactions within populations. Our findings shed light how SAC could operate at the molecular level, and sets a standard for future investigations.

## Introduction

Sexual conflict arises when the fitness optima of a trait shared by the sexes are not aligned (Parker, 1970, 1979; Arnqvist & Rowe, 2005; Perry & Rowe, 2015). Interlocus sexual conflict occurs when different loci in males and females influence the phenotypic value of a trait that depends on contributions from both partners. A classic trait influenced by interlocus sexual conflict is the female remating rate (Parker, 1970, 2020). Males benefit when they can prevent future inseminations by rival males, for example by reducing female receptivity to further matings. In contrast, females may gain fitness, either *via* direct or indirect benefits, from controlling when, how often, and with whom to remate (Arnqvist & Nilsson, 2000; Rundle et al., 2007; Priest, Galloway, et al., 2008; Priest, Roach, et al., 2008; Fromonteil et al., 2023). There can also be strong selection on males to court females aggressively, even if they are likely to have already been mated and therefore unreceptive, if the opportunity exists for them to accrue benefits from such courting, and thereby increase the overall female mating rate (Arnqvist & Rowe, 2002a; Perry & Rowe, 2012). Such conflicts can result in sexually antagonistic co-evolution (SAC), an escalating arms race between the sexes leading to novel and increasingly exaggerated persistence and resistance traits in males and females (Holland & Rice, 1998; Arnqvist & Rowe, 2002a, 2005; Perry & Rowe, 2015). On average, males and females should be evenly matched in this arms race, making conflicts difficult to detect (Arnqvist & Rowe, 2002a). However, a central prediction of SAC theory is that due to the cyclical nature of SAC, transient mismatches can occasionally result wherein for some populations (or species) either males or females have the upper-hand, that is, the value of the shared trait (e.g. the female remating rate) is more closely aligned with the optimum for one sex. Which sex has the advantage can be detected from patterns of relative phenotypic exaggeration of male and female traits (Arnqvist & Rowe, 2002a, 2005).

The most well-documented example of SAC influencing phenotypic traits is among *Gerris* water striders (Arnqvist & Rowe, 2002a, 2002b). Intense struggles between the sexes over the mating rate, an inherently shared trait, generates strong selection for male persistence and female resistance morphologies and behaviours (Arnqvist & Rowe, 2002a, 2002b). In their landmark study of SAC in this system, Arnqvist & Rowe (2002b) quantified the co-evolution between male and female traits that mediate the conflict using species-level measures of average male and female morphologies. Using 2-Block Partial Least Squares (2B-PLS) they found that male and female morphologies, represented by matrices of species-level mean morphometric landmark coordinates, show significant co-variation across species (Arnqvist & Rowe, 2002a). Furthermore, principal component analysis (PCA) of the main axes of co-variation between male and female traits identified two principal components (PCs), where PC1 reflects the main axis of co-variation, while PC2 reflects the species-level *relative* difference or exaggeration, (“mismatch”, or “imbalance” in Arnqvist and Rowe 2002a) between the sexes within a given species (Arnqvist & Rowe, 2002a). In their analysis, a higher value on PC2 was associated with a greater exaggeration of female resistance traits than male persistence traits, which the authors used to infer a female advantage. Conversely, a lower value on PC2 was used to infer a male advantage. If SAC drives the observed co-variation in these traits, then PC2 should therefore predict the outcomes of mating interactions within each species. Populations with inferred female advantage populations should have values of shared traits under sexual conflict (e.g. length of pre-mating struggles and male mating success) closer to the female optimum. Perry & Rowe (2012) have used the same approach to quantify co-variation across populations of the waterstrider *Gerris incognitus*. Across both species and populations, the relative level of exaggeration between male and female morphologies did in fact predict the outcome of mating interactions, reflecting which sex has an advantage, in agreement with SAC theory (Arnqvist & Rowe, 2002a; Perry & Rowe, 2012). Relatively more exaggerated male morphologies resulted in, for example, shorter pre-mating struggles, and higher male mating success (Arnqvist & Rowe, 2002a; Perry & Rowe, 2012). Patterns of co-variation in antagonistic male and female traits in other taxa also likely reflect SAC (Bergsten et al., 2001; Bergsten & Miller, 2007; Rönn et al., 2007; Tatarnic & Cassis, 2010; Perry & Rowe, 2015), making this a compelling hypothesis explaining phenotypic diversity across several systems.

As genetic evolution underlies phenotypic evolution, the logic of SAC should also extend to the genetic level at loci underlying male and female traits that mediate sexual conflicts. Alleles at genes underlying these traits interact via the phenotypes that they produce and different alleles of male persistence traits will then be better or worse matches, with respect to the male optimum value of the shared trait, against alleles underlying female resistance traits. Thus, these loci (e.g. Swanson & Vacquier, 2002, and reviewed in Sirot et al., 2015; Rowe et al., 2018) should show patterns of co-varying differentiation and divergence between male- and female- beneficial loci across populations and species. Indeed, genetic (co-)differentiation, meaning population-level differences in allele frequencies, of male and female traits and their underlying genes is an outcome of population genetic models of SAC (Frank, 2000; Gavrilets & Waxman, 2002; Hayashi et al., 2007). Such patterns of differentiation would reflect population- or species-level differences in the phenotypes underpinned by these loci.

Under SAC theory, relative differences in the degree of population genetic differentiation at loci underlying male- and female- traits should therefore predict which sex has the upper-hand and fitness-related outcomes of mating interactions, by the same logic as for phenotypic traits (Arnqvist & Rowe, 2002a, 2002b; Perry & Rowe, 2012). However, no studies have tested these predictions, largely because the loci underlying most conflict traits remain unknown (but see Khila et al., 2009; Crumière & Khila, 2019; Pruvôt et al., 2025). Meanwhile, classes of loci proposed to be associated with sexual conflicts and SAC, such as genes encoding male seminal fluid proteins (SFPs), show rapid evolutionary change across species (Swanson & Vacquier, 2002; Haerty et al., 2007). These SFPs, transferred to females during mating, mediate female reproductive physiology, with effects on female traits such as receptivity to further mating attempts, egg production, and survival (Sirot et al., 2015). While SFPs are relatively well-described genetically with many having known phenotypic functions, their co-evolution with loci underlying female resistance traits is unexplored and how genetic (co-)variation at these loci impacts fitness outcomes of male-female interactions is unknown. Thus, unlike the progress of the “genomics revolution” in understanding the genetics of ecological adaptations over the past quarter century, studies of the genomics of sexually selected and sexual conflict traits have lagged behind (Wilkinson et al., 2015; Kasimatis et al., 2017; Rowe et al., 2018).

Empirical tests of SAC theory require integrating genetic (co-)variation at known SAC loci and the consequence of the co-variation for outcomes of mating interactions across multiple populations and species. Such tests are critical for understanding several facets of the genetics of SAC; how many loci, and what proportion of those underlying a given trait, are involved in this ubiquitous process in sexually reproducing taxa? How do sex-specific selection and co-evolution between males and females shapes patterns of genetic variation across populations and species? And what are fitness consequences of this genetic (co-)variation (Wilkinson et al., 2015; Kasimatis et al., 2017; Rowe et al., 2018)?

We capitalize on the iconic “sex peptide” (SP) network of *Drosophila melanogaster* to explicitly test SAC theory in a molecular framework for the first time. The SP-network includes a male component, the SP protein and a suite of other SFPs, that interact with a female component, “sex-peptide receptor” (SPR) and other proteins expressed in the female reproductive tract (FRT), although we note that several of these loci are expressed to varying degrees in both sexes (Kubli & Bopp, 2012; Findlay et al., 2014; Singh et al., 2018; Hopkins & Perry, 2022; see also below). Through these interactions, SP has been shown to influence a wide range of female post-mating behaviours. For example, it depresses the female remating rate, induces a higher ovulation rate, shortens female lifespan, and reduces reproductive success (reviewed in Kubli & Bopp, 2012; Hopkins & Perry, 2022). These traits are classic sexual conflict traits and may therefore also potentially subject to SAC in *D. melanogaster* (Parker, 1970; Rundle et al., 2007; Priest, Galloway, et al., 2008; Priest, Roach, et al., 2008; Smith et al., 2017; Parker, 2020). However, whether the evolution of *SP* and *SPR* is driven by SAC is controversial (Hopkins & Perry, 2022; Hopkins et al., 2024). Controversy arises because evidence that SP transfer results in reductions in female lifespan or lifetime reproductive success is mixed (Hopkins & Perry, 2022). Additionally, across species, molecular evolution of *SP* and *SPR* seems uncorrelated (Findlay et al., 2014; Hopkins et al., 2024) contrary to the predictions of SAC theory (Parker, 1970; Arnqvist & Rowe, 2005). However, other traits influenced by SP, such as the female remating rate and oviposition patterns, may be equally important sources of sexual conflict in natural populations (e.g. Wensing & Fricke, 2018). These traits are frequently ignored in assessment of the role of sexual conflict and SAC in SP and SPR co-evolution (but see e.g. Kohlmeier et al., 2021). Additionally, while SP and SPR interact directly as ligand and receptor, the other components of the SP-network may interact more indirectly. Indeed, correlations in the evolutionary rate of these other SP-network loci have been observed (Findlay et al., 2014), yet these other loci of the male- and female- components of the SP-network are also frequently ignored.

If SAC drives the evolution of the loci of the SP-network across populations and species, then, like SAC on morphological traits, we expect to find co-variation in genetic differentiation and divergence of these loci. This is expected regardless of the mode of evolution; differentiation and population divergence could be due to the origin and sweep to fixation of novel mutations, or it could follow a frequency-dependent process where sets of alleles at some loci fluctuate in frequency over time, in response to the frequency of antagonistic alleles at other loci, or a combination of the two. Moreover, while there is genetic variation within the coding sequence of SP, that may be functionally related to the interaction with SPR, SP gene expression differences between inbred lines are also associated with variation in outcomes of mating interactions (e.g. remating rates; Smith et al., 2009). Thus, there is no *a priori* reason to expect that coding sequence variation should be more important than regulatory sequence variation driving gene expression differences in this system. Any measures of population differences at these loci must be equally agnostic in these respects.

The above features of a locus are conveniently captured by the measure of genetic differentiation, Fst, which is an estimate of differences in allele frequencies between populations (Weir & Goudet, 2017; Hivert et al., 2018). Thus, although quantifying the co-variation in levels of differentiation across the loci of the SP-network does not necessarily indicate functional divergence between populations, because precise causal variants are unknown, such a pattern is expected from SAC. Moreover, we can make the further prediction that if such a pattern is driven by SAC, then, similar to SAC on morphological traits, relatively more genetic differentiation in the male- or female- component should associate with outcomes of mating interactions closer to the male- or female- optimal trait value, respectively. Thus, relative differences in population-level allele frequencies, measured as genetic differentiation, should reflect a relative degree of “molecular exaggeration” of loci, and the underlying phenotypes. This genetic measure is therefore analogous to the relative differences between the sexes in mean population- or species-level morphologies of water striders in the canonical study of Arnqvist & Rowe (2002a) and others (Perry and Rowe 2012).

Leveraging the known loci of the male- and female- components of the SP-network, we use population genomic data across ∼150 natural populations of *D. melanogaster* (Kapun et al., 2021; Nunez et al., 2025), combined with lab populations established from a resource of isofemale lines of natural populations (Durmaz Mitchell et al., 2025), to evaluate the extent and consequences of co-evolutionary patterns within and between the male- and female- components of the SP-network, using methods that parallel morphological studies of SAC, to test predictions of SAC theory at the molecular level for the first time.

## Methods

### Genetic data

Population genomic data were obtained from the publicly available DEST dataset which comprises a total of 530 pool-seq samples of ∼46,000 flies from >170 populations world-wide sampled across years (https://dest.bio/; (Kapun et al., 2021; Nunez et al., 2025). Using hierarchical Fst analysis of a neutral set of SNPs with poolfstat (Hivert et al., 2018; Gautier et al., 2024; see “1.1. Hierarchichal Fst analysis” in the Supplementary Materials), we found that samples from the same population showed overall low within-population Fst (Fsg = 0.01, [0.0096, 0.0104]), compared to between-population Fst (Fgt = 0.04 [0.038, 0.045]), in agreement with previous results (Nunez et al., 2025). Thus, we chose a single pool-seq sample per population for further analysis, favouring samples with highest coverage, to ensure the highest quality data and estimates. Populations from the ancestral range in Africa were excluded due to very few samples, different sequencing strategies, and a large impact on overall diversity and differentiation so that the final set of samples represented 149 populations (Table S1).

A set of interacting loci that make up the SP-network have been described from several studies. Here we use the union of loci described in Findlay et al., (2014) and Singh et al., (2018), because these represent the current most complete understanding of the system (see also McGeary & Findlay, 2020). Several loci are poorly characterised in this context, but all have been shown to be necessary for the observation of SP-network mediated post-mating behaviours (in particular the reduced female receptivity to remating; Findlay et al. 2014).

These loci can be divided into “male”- and “female”- components of the SP-network based on sex-specific reproductive tissue expression (Tables S2 and S3) although this does not necessarily preclude expression of those loci in the other sex. For example, SPR is expressed only in the female reproductive tract (FRT) but is expressed in the nervous system of both sexes (Table S3). We consider two different definitions of the SP-network loci, depending on the analysis. Where only coding sites are analysed (πN/πS), we limit ourselves to the coding sequence (CDS) of each gene. In other analyses (Fst, π, and Tajima’s D) we include introns and sites 2,000 bp up-/down-stream of the locus. This is to strike a balance between a focus on the loci of interest while ensuring enough data for accurate population genetic estimates and inclusion of potential regulatory elements. To obtain genomic background estimates as a comparison, we also sampled 100 sets of 13 randomly selected loci (1,247 unique loci, ∼9% of the annotated protein-coding loci) from the *D. melanogaster* annotation.

To investigate patterns of genetic diversity and evaluate the role of selection at SP-network loci, population genetic summary statistics were computed (π, Tajima’s D, and πN/πS), with npstat (v. 1.0; Ferretti et al., 2013) and SNPGenie (v. 2019.10.31; Nelson et al., 2015). For πN/πS, we summed the mean number of pairwise (non-)synonymous differences and the number of (non-)synonymous sites, across all populations for each gene. We then computed overall πN/πS for each locus using these sums. To derive confidence intervals we bootstrap sampled populations with replacement and re-computed πN/πS 100 times. We also computed population-specific Fst values (see “1.2. Definition of population-specific Fst” in the Supplementary Materials) for each locus in the SP-network with poolfstat (v. 2.1.2; Hivert et al., 2018).

### Analysis of molecular co-evolution

Population-specific Fst estimates were analysed to characterise patterns of co-variation in Fst between male- and female- components of the SP-network. We use 2-Block Partial Least Squares (2B-PLS) to identify male- and female- axes that describe the largest amount of co-variance in Fst between male- and female- components of the SP-network. 2B-PLS is well suited for analyses of co-evolution across species or populations (Rohlf & Corti, 2000) and has been employed to study complex multivariate morphologies in the context of SAC (e.g. Arnqvist & Rowe, 2002a, 2002b; Perry & Rowe, 2012). 2B-PLS produces two main reduced axes that describe co-variation between the two blocks, as well as individual loading scores for the components of those blocks on the axes. A permutation test of rows within the two blocks tests whether the degree of co-variation is greater than expected by chance. This permutation test conserves relationships between the variables (in this case the loci of the SP-network) within blocks, but should break any signal of co-variation across blocks.

Populations are not independent but genetically related to varying degrees by shared evolutionary history and gene-flow. Such population structure could produce correlations between loci, and should therefore be accounted for in analyses (Stone et al., 2011; Castillo, 2017; Chelini et al., 2021; Raas & Dutheil, 2024). 2B-PLS also integrates phylogenetic or relatedness information using a phylogenetic tree (Adams & Felice, 2014). We first computed the omega matrix, a relatedness matrix, for all populations using BayPass (v. 2.4; (Gautier et al., 2013); see “1.3. Computation of the relatedness matrix” in the Supplementary Materials) and then converted the summarised omega matrix of BayPass to a dendrogram using the neighbour-joining method in the “ape” R package (v. 5.8; (Paradis et al., 2004). This relatedness matrix accounts for any broader population-genomic structure which should affect the whole genome equally. Phylogenetically corrected 2B-PLS was used to test for and describe the main axes of co-variation between male- and female- components of the SP-network with the “geomorph” R package (v. 4.0.8; Adams et al., n.d.; Baken et al., 2021). Because the sign of the strongest loadings on both axes was negative (Table S4), we switched the sign of all values on both axes for ease of interpretation and plotting, such that higher values on the 2B-PLS axes indicated higher differentiation at the loci with strongest loadings on these axes. The main axes of co-variation from the 2B-PLS analysis can be further processed into two principle components by PCA, where the first PC describes the absolute degree of genetic differentiation across both male- and female- components of the SP-network. Meanwhile, the second PC axis describes the *relative* differences between the sex-specific components of the SP-network in the degree of genetic differentiation and provides a continuous measure of the population-level degree of genetic “mismatch” or “molecular exaggeration” between the sexes, as for morphological SAC traits (Arnqvist & Rowe, 2002a, 2002b).

To complement the internal permutation test and phylogenetic control, we also conduct a bootstrapping procedure to investigate patterns of co-variation in population genetic differentiation and the identification of outlier populations using randomly selected sets of loci. For each of the 100 sets of 13 loci sampled above (see “Genetic data” above) we computed all pairwise correlations, and conducted 2B-PLS exactly as for the SP-network loci (randomly assigning genes to a “male” or “female” block in the same 9:4 proportions as for the SP-network loci). We then computed the principle components of the male- and female- axes of co-variation from 2B-PLS analysis (see above) and compared the rank order of populations along PC2 from each random set of loci to that observed for the SP-network using Spearman rank correlation tests.

### Behavioural experiments

We set up stock population cages using isofemale line resources available through the DrosEU consortium (Durmaz Mitchell et al., 2025; see “1.4. Isofemale line resources and stock population cages” in the Supplementary Materials). Briefly, we created three replicate population cages using isofemale lines from each of six natural populations (18 cages in total). For each population, we took 15 males and 15 females of each isofemale line, and split them across the three replicate cages. Populations were allowed to mate freely and recombine for 4-9 generations prior to the experiment. Experimental flies were obtained from each population cage using egg laying plates consisting of a molasses, agar, and water mix (in 1L dH_2_0: 16.7g agar, 159.7g molasses) set in a small Petri dish lid (4 cm in diameter) and topped with a small amount of yeast paste (in 10mL dH_2_0: 3g dried yeast) placed in each cage overnight. 100 eggs from each cage were collected and distributed across 5 vials (20 eggs / vial) containing standard fly medium (see Supplementary Materials). These were left to develop for 9-11 days in an incubator (Panasonic MIR-154-PE, Panasonic Healthcare Co. Ltd., 25° C, 12h:12h L:D). On the days of eclosion, virgin flies were collected from each vial throughout the day, ensuring collected flies were no older than 2 hours post-eclosion and thus virgin. Flies were kept in single-sex groups of up to 10 flies in vials containing standard fly medium for 3 days prior to mating and remating assays. Collections of eggs and virgins for mating and remating assays were staggered in order to obtain, as far as possible, males of controlled ages. All males were 2-5 days old at the time of mating and remating assays.

On the day of the mating assay, a female and male from the same population were loaded into individual vials. We recorded the time at which pairs entered the vial, and both the start and end time of copulation. The assay was run for 2 hours from the time of the last pair entering the vial. Flies that did not mate in that time were discarded (∼3-12% of all flies across populations). There was no relationship between the percentage of “failed” pairings and the relative degree of genetic differentiation (Pearson’s correlation: cor = −0.17, t = −0.34, d.f. = 4, p = 0.75). Flies that were observed to mate later than 2 hours after pair formation were censored and discarded. Mated males were removed from their vial and females were left to lay eggs for 2 days. A female remating assay was performed the second day after the mating assay in which each female was transferred to a new vial and paired with a new male from the same population of approximately equal age to the initial male to isolate the effect of remating. We again recorded the time at which pairs entered the vial, and, if remated, copulation start and end times. To account for female remating refraction, we ran the assay for 3 hours from the time of the last pair entering the vial, pairs that were observed to mate later than 3 hours after pair formation were censored and discarded.

We collected data on two fitness related traits for each female – mortality and offspring production. Mortality was assessed for each female up to 30 days after the first mating day. All vials were checked every day (except weekends) and the date of death recorded for each female. Females that died due to handling error or escaped were censored as were all females that survived past the 30-day cut-off. Throughout the experiment, females were provided with fresh vials of fly medium on days 2, 4, 7, 11, 17, and 24 after the mating assay, producing 7 vials of offspring in total. Total reproductive success was scored by counting offspring from each vial after 14 days, to allow ample time for the majority of offspring to hatch, pupate, and eclose as adults. Total reproductive success, was measured as either the total offspring in vials 2 and 3, or the total offspring across all 7 vials. This was done to keep a maximum amount of data, as some females died during the experiment and did not have an opportunity to lay eggs in later vials. We also measured timing of peak offspring production computing the time, in days, until the vial with the maximum number of offspring. The full experiment was conducted over five batches to obtain suitable sample sizes and batch was included as a random factor in analyses.

In a second mortality experiment, experimental flies were collected from each cage using egg laying plates as above. Upon collection of emerged adults, virgin flies were distributed across vials in groups of 10 females and 10 males per vial. Vials were then kept in a controlled temperature room (25° C, 12h:12h L:D) for the duration of the experiment. Survival of females was checked every day, except weekends, until less than 10% of flies from each cage were remaining.

See the Supplementary Materials for the details of all statistical analyses.

## Results and Discussion

To assess molecular evidence for SAC in the male- and female- components of the SP-network across *D. melanogaster* populations we first evaluated the overall population genetic signature of selection at the loci of the SP-network. Comparative work has found rapid evolution of *Drosophila* SFPs generally, and co-variation in the evolutionary rates across species among the loci of the SP-network in particular (Haerty et al., 2007; Findlay et al., 2014; Hopkins et al., 2024). However, *SP* and *SPR* seem to have uncoupled evolutionary rates across species (Findlay et al., 2014; Hopkins et al., 2024). These patterns are consistent with either relaxed- or positive-selection. Selection (whether positive, in favour of new beneficial alleles, or purifying, against new deleterious mutations) should leave a characteristic signal of reduced non-synonymous diversity (πN) relative to synonymous diversity (πS), comparable to other protein-coding genes subject to selection. We find that neither nucleotide diversity (π) nor Tajima’s D for SP-network loci are different than for the set of randomly selected loci (Figure 1A), and πN/πS values are well below 1 (Figure 1A), patterns of strong evidence of selection. Combined with the observed rapid evolution across species at SP and other SFPs (Haerty et al., 2007; Findlay et al., 2014; Hopkins et al., 2024), the sum of the evidence, is consistent with a process of recurrent selective sweeps. Such a dynamic genomic signature is expected for SAC traits, in which alleles beneficial to one sex rapidly spread, but which are then countered by new alleles beneficial to the other sex.

**Figure 1.**
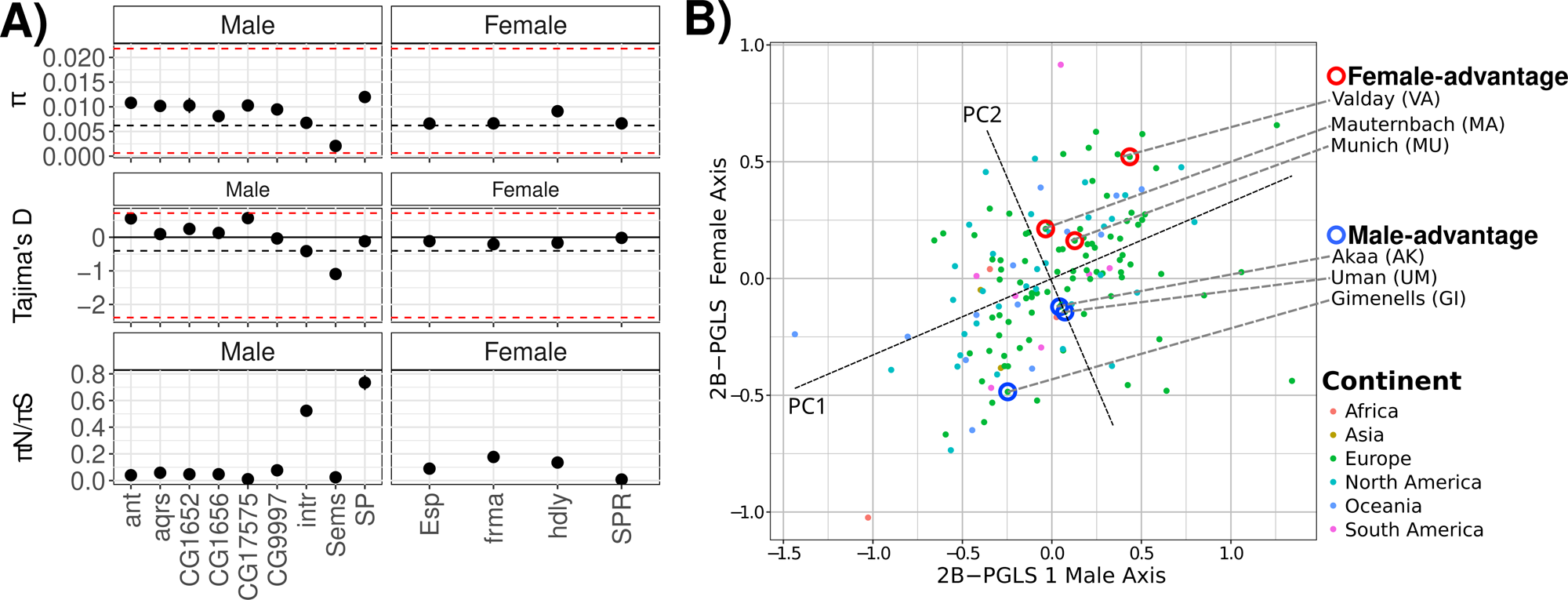
**A)**. Estimates of π (top), Tajima’s D (middle), and πN/πS (bottom) for loci in the male- and female-components of the SP-network. Red dashed lines show the maximum and minimum values obtained for a set of 100 randomly sampled genes from the rest of the genome. **B)** Male and female axes of co-variation in Fst at SP-network loci. Points are coloured by their continent of origin. Populations chosen for behavioural experiments, and the sex with inferred advantage are highlighted with coloured circles. The dashed lines labelled “PC1” and “PC2” illustrate the first two principle components from a PCA of the male and female axes of co-variation.

We next found strong evidence for co-variation in population genetic differentiation between the male- and female- components of the SP-network from analysis of population-specific Fst estimates of each locus (Phylogenetically corrected 2-Block Partial Least Squares (2B-PLS) analysis; D. C. Adams & Felice, 2014; Rohlf & Corti, 2000; r-PLS = 0.44, Z = 4.17, p = 0.001; Figure 1B).

The male axis of this co-variation loads most heavily for SP network genes, *SP* and *Sems*, while the female axis loads most heavily for *Esp* and *hdly* (Table S4). There is relatively low loading of *SPR* on the female axis, which is consistent with the weak patterns of covariation between *SP* and *SPR* observed across species (Findlay et al., 2014; Hopkins et al., 2024; Table S4). In summary, across 149 natural *D. melanogaster* populations, our results are consistent with ongoing selection at SP-network loci, and natural genetic co-evolution between male- and female- components of the network are in line with predictions from SAC theory. The pattern of co-variation is weak between the most directly interacting *SP* and *SPR*, as has been found in comparative results from co-evolutionary analyses of these loci (Hopkins et al., 2024; McGeary & Findlay, 2020). We find the strongest evidence for co-variation between *SP*, *Sems* and other indirectly interacting loci of the female component of the SP-network (*Esp* and *hdly*). These also show the highest correlations in Fst in simple pairwise tests (Figure S1). Our results highlight that the most obvious candidate loci underlying conflict traits may not always be the strongest drivers of SAC, and that only a subset of the loci underlying a suite of traits mediating sexual conflicts, such as the SP-network, may be active in SAC. By including multiple interacting loci of SP-network, rather than focusing on the most intuitive one from each sex, we are able to detect molecular signatures of co-evolution that may otherwise be elusive.

A central prediction of SAC is that relatively more exaggeration in male- compared to female- traits infer a fitness advantage in sexual conflicts to males and *vice versa* (Arnqvist & Rowe, 2002a, 2005). We extend this prediction to the molecular level, defining the relative difference in genetic differentiation between male- and female- components of the SP-network as a population-level molecular “mismatch” or “exaggeration”, expressed as a continuous measure using the second principle component (PC2) of the male- and female- axes of co-variation derived from the 2B-PLS – precisely as has been done for morphology traits (Arnqvist & Rowe, 2002a, 2002b; Perry & Rowe, 2012; Figure 1B; Table S5). Higher scores indicate greater genetic differentiation in allele frequencies from the overall average, in the male- compared to the female- component and an inferred advantage to males. If these patterns of genetic co-variation between components of the SP-network, with previously described phenotypic effects (Kubli & Bopp, 2012, but see Hopkins & Perry, 2022), reflect a process of SAC, then populations with an inferred male- advantage should have: lower female remating rates and an earlier peak in offspring production (faster initial rate of offspring production) than populations with an inferred female-advantage. If SP harms females by reducing lifespan, then we also expect shorter total lifespan and lower survival in populations with inferred male advantage.

We tested these predictions using six populations along the axis of relative genetic differentiation (PC2; Figure 1B), representing a continuum from strong inferred male-advantage to strong inferred female-advantage, for which isofemale line resources are available; these represent a small proportion of the original 149 populations (Durmaz Mitchell et al., 2025; Figure 1B, Table S5). As our aim was to investigate population-level phenotypic effects, we reconstructed each of the six populations by combining individuals from all isofemale lines for each population in population cages, with three replicate cages per population, and assayed female remating rates, survival, offspring production per vial, and total offspring number (see Methods and Supplementary Materials).

Remating rates varied across populations, from 14% (Uman, Ukraine) to 40% (Munich, Germany) (Figure 2A; Table S6; total N = 765, n= 113-140 females per population). Critically, the variation in remating rates confirmed the predictions from SAC theory: populations with relatively more genetic differentiation in the male component of the SP-network, i.e. greater inferred male-advantage, had lower female remating rates (Figure 2A and 2B). Over 94% of the posterior distribution of the relationship between relative genetic differentiation and female remating rates was < 0, indicating a clearly negative relationship, as predicted by SAC theory (Figure 2B). The model in which relative genetic differentiation (PC2) as the explanatory variable was represented by the degree of male- and female- advantage was preferred over a model that did not include this explanatory variable (ELPD_w/ term_ = −399.3 ±14.7, ELPD_w/o term_ = −399.4 ±14.6, ELPD difference = −0.1 ±0.7). These effects are even stronger when contrasting female remating rates from populations with the greatest inferred male- and female- advantage, with 99% of the posterior distribution supporting a lower remating rate for populations with an inferred male advantage (Figure S2).

**Figure 2.**
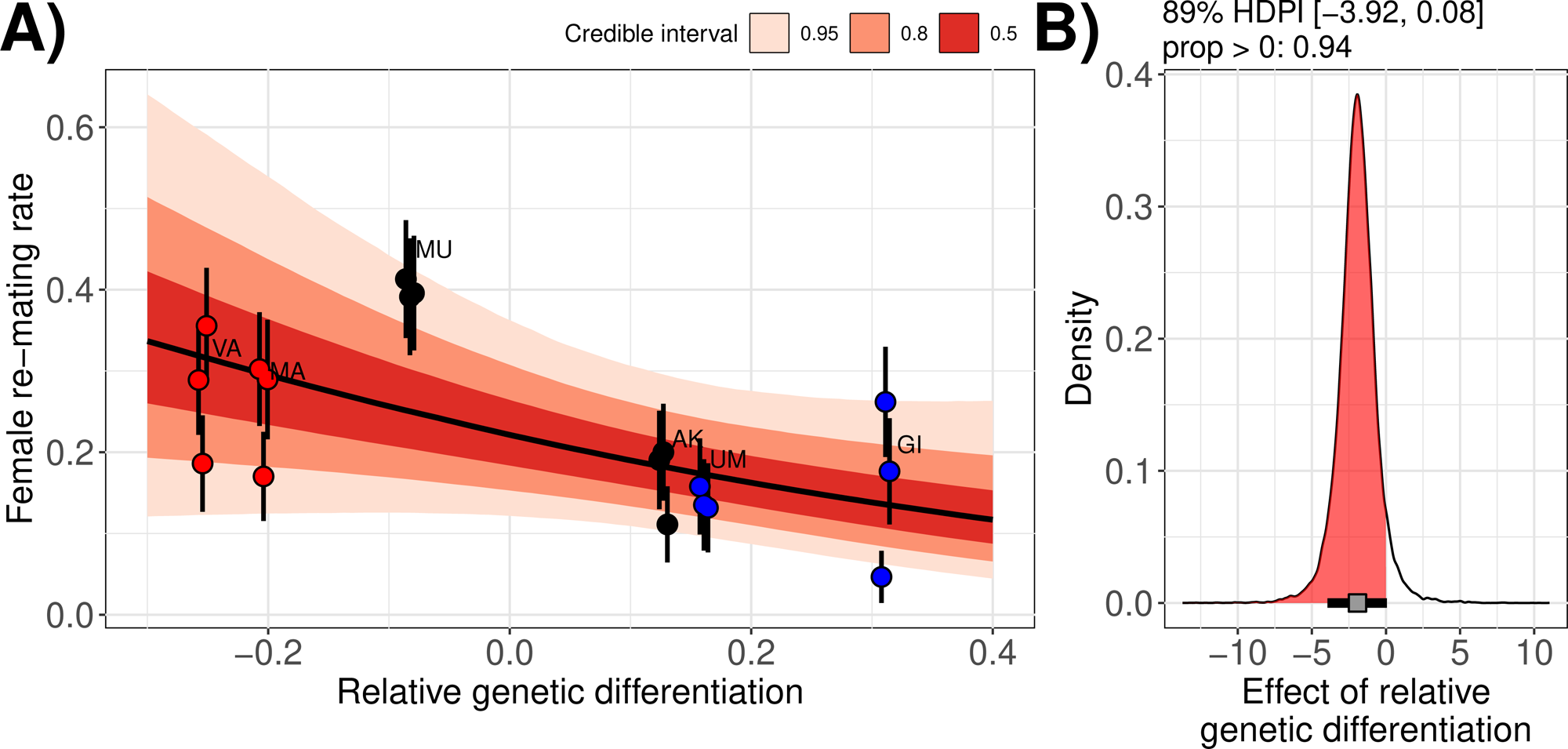
**A)** Female remating rates as a function of a continuous measure of relative genetic differentiation at female- and male- components of the SP-network. Negative values indicate relatively more genetic differentiation in the female- versus the male- component. Each individual point is a replicate population cage, points are coloured by which sex is inferred to have an advantage in sexual conflicts (red – female-advantage, blue – male advantage, black – intermediate). The slope and error ribbons represent the posterior mean and credible intervals for the slope. **B)** Posterior distribution of the slope estimate. The area shaded in red indicates the proportion of the posterior distribution with slope estimates < 0, that supports a negative relationship between relative genetic differentiation and the female remating rate, the grey box and bar indicate the posterior mean and 89% HDPI respectively.

The relative difference in genetic differentiation between the male- and female- components of the SP-network was also associated with patterns of oviposition, as predicted by SAC theory. Populations with an inferred male advantage had peak oviposition earlier than populations with an inferred female advantage (Figure 3; Figure S3). This resulted in females from populations with inferred female advantage producing 68% of their total offspring within the first 7 days after mating, on average, compared to 71% for females from populations with inferred male advantage. Male induction of faster egg-laying rates and earlier peaks of offspring production bias paternity to the most recently mated male and may be counter to female interests insofar as they override female-beneficial determination of the timing of reproductive investment (i.e. cryptic female choice; Eberhard, 1996; Arnqvist & Rowe, 2005). Indeed, experimental evolution studies that manipulate the opportunity for sexual selection and conflict have found that both male ability to reduce female remating rates (Crudgington et al., 2005) and oviposition timing (Crudgington et al., 2005; Edward et al., 2010) readily evolve in response to these treatments, strongly suggesting that these traits are under sexual conflict.

**Figure 3.**
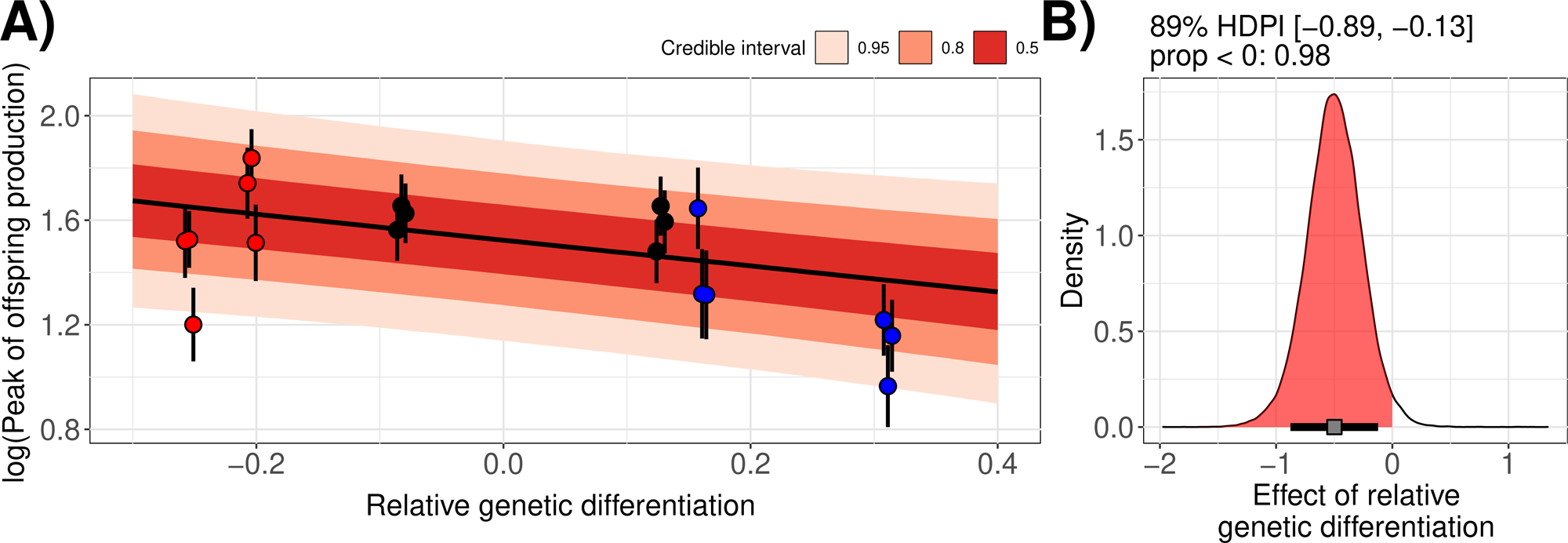
**A)** The log(number of days) until the peak of offspring production as a function of a continuous measure of relative genetic differentiation at female- and male- components of the SP-network. Negative values indicate relatively more genetic differentiation in the female- *versus* the male-component. Each individual point is a replicate population cage, points are coloured by which sex is inferred to have an advantage in sexual conflicts (red – female-advantage, blue – male advantage, black - intermediate). The slope and error ribbons represent the posterior mean and credible intervals for the slope. **B)** Posterior distribution of the slope estimate. The area shaded in red indicates the proportion of the posterior distribution with slope estimates < 0, that supports a negative relationship between relative genetic differentiation and the peak of offspring production, the grey box and bar indicate the posterior mean and 89% HDPI respectively.

Contrary to SAC predictions, we found no relationship between the relative difference in genetic differentiation between the male- and female- components of the SP-network and total number of offspring (Figure S4). Similarly, while there is strong evidence that survival differs between populations (Cox regression survival analysis; X^2^ = 27.03, d.f. = 5, p < 0.001; Figure S5A), there was no evidence that this varied based on which sex was inferred to have an advantage (HR = 1.23, Z-ratio = 1.04, p = 0.3). Moreover, there was no evidence that either female remating status (X^2^ = 0.36, d.f. = 1, p = 0.55) or the interaction between remating status and population (X^2^ = 4.66, d.f. = 5, p = 0.46) influenced survival. Survival results were similar in the second experiment tracking survival over ∼80 days, while also allowing for continuous mating in groups of 10 males and 10 females (Figure S5B).

Using a bootstrapping approach, we found that within sampled sets of 13 loci, mean significant pairwise correlation coefficients for population-specific Fst were greater than for the SP-network in only 14% of cases. Additionally, positive rank correlations of values on PC2 for populations for which isofemale line resources exist between the random sets of loci and those of the SP-network loci was found in only 12% of cases, never exceeded correlation coefficients of 0.9, and never passing stringent Bonferroni correction for multiple testing. Thus, despite strong evidence for co-variation in population-specific Fst recovered for many sets of randomly selected loci (78% with p < 0.001), the pattern of populations that predicts female re-mating rates is not one which is common throughout the genome and instead reflects patterns of co-variation more specifically between the loci of the SP-network. Evidence for co-variation at randomly selected loci might be explained by, for example, concerted evolution due to selection for conserved interactions between proteins, or demographic processes not fully captured by the relatedness matrix.

By using patterns of natural genetic (co-)variation of loci of multiple male- and female- SP network components across multiple populations, we explicitly test predictions made by SAC and find that our results are consistent with this process. Interest in *Drosophila* SFPs as mediators of sexual conflict arose because of their apparent toxic effects to females, observed as reduced lifetime reproductive success and survival (Kubli & Bopp, 2012; Hopkins & Perry, 2022). However, the role of sexual conflict in the (co-)evolution of SP and SPR has been questioned (Kubli & Bopp, 2012; Hopkins & Perry, 2022). This is due mainly because of inconsistent results across studies for the expected reductions in female longevity or lifetime reproductive success, leading to speculation that these costs may be highly condition or environment dependent, or that these traits are not sexually antagonistic at all (Hopkins & Perry, 2022). We found no relationship between this exaggeration on total offspring production or lifespan, suggesting that these traits, in fact, are not driven by SAC at these loci. However, fewer studies have focussed on female remating rates and the timing of offspring production, despite these traits potentially also being under sexual conflict and highly relevant in nature (Eberhard, 1996; Arnqvist & Nilsson, 2000; Arnqvist & Rowe, 2005). We show that both the female remating rate and egg production timing are predictable based on relative differences in genetic differentiation between male- and female- components of the SP-network. Thus, our results extend SAC theory to the molecular level for the first time.

However, the extent of the role of conflict in the phenotypes mediated by the SP-network remains unclear. Full accounting of the fitness costs and benefits to females is difficult. Therefore, just as with the mixed evidence for survival costs of SP transfer to females (reviewed in Hopkins & Perry, 2022), there is ambiguity in the balance of costs and benefits to females from other post-mating traits. For example, Kohlmeier et al. (Kohlmeier et al., 2021) present evidence that reduced receptivity to female remating may be due to increased choosiness, mediated by reduced sensitivity of female olfactory neurons to male pheromones of mated females compared to virgins. Kohlmeier et al. (2021) show that this effect is reproducible also on injection of SP. How this elevated choosiness translates to benefits is unclear especially given that SP not only reduces receptiveness to rematings but also increases egg-laying rates. Moreover, even increased post-mating choosiness can be interpreted as an outcome of sexual conflict rather than sexual selection. Theoretical modelling has also shown that trading off late-life reproduction in favour of early-life reproduction can be beneficial to females if they are adapted to or experience otherwise high extrinsic mortality and are unlikely to reap the benefits of delayed egg-laying (Bonduriansky, 2014). Thus, even if post-mating egg-laying rates are under sexual conflict, in some circumstances the costs to females may be low or it may even be beneficial. In general, environmental and ecological variables can dramatically modulate the effects of sexual selection and conflict (García-Roa et al., 2020). This raises the possibility that the role of conflict might vary across populations and with ecological variables, complicating the dynamics of SAC which would require taking these ecological variables, and their effects, into account. Indeed, in the classic water-strider system, studies have shown that ecological variables also explain morphological variation mediating mating struggles (Perry et al., 2017). Thus, a straightforward role for sexual conflict in these phenotypes remains elusive, and more research placing the outcomes of SP-mediated effects in an ecological context are critically needed.

Nevertheless, our study addresses critical shortcomings in previous work and offers an explanation to seemingly contradictory results. SAC theory predicts that sex-specific effects will be likely be highly dependent on genotype-by-genotype interactions (Pizzari & Snook, 2003; Rowe et al., 2003; Wensing & Fricke, 2018), so mixed results across studies using either different populations or inter-population crosses is unsurprising and actually expected (e.g. Smith et al., 2009; Wensing & Fricke, 2018). Our study, by taking an explicitly comparative approach and using within-population crosses, avoids this unpredictability, and instead focusses on patterns of co-evolution between males and females, across populations. Additionally, our study also highlights the perils of a specific focus on genetic variation in, and the direct interactions of, only *SP* and *SPR;* the SP-network consists of several other male and female proteins that interact more indirectly but may play a more important role in the co-evolution between males and females. We show that effects of molecular exaggeration at SPR is relatively small and that, instead, other loci of the female component of the SP-network is driving female resistance evolution. Not accounting for the effects of variation at these loci will miss much of the relevant biology.

## Conclusions

In sum, we extend SAC to the molecular level for the first time. We predict and test outcomes of mating interactions by documenting patterns of natural genetic co-evolution and genetic differentiation at interacting loci of the classic *Drosophila* SP-network. Our results show that relatively few interacting loci can drive SAC, and predict sexually antagonistic phenotypes, which, coupled with strong selection, may explain the rapid pace of this evolutionary process, at least for some traits and genes. This also offers encouragement to other studies of sexually antagonistic complex behavioural and morphological traits in that, while good knowledge of the relevant interacting loci is needed, not all loci underlying these traits need be accounted for. Our approach offers a standard that can be extended to ongoing work identifying loci underlying conflict traits in other systems to yield important insights into how SAC can shape patterns of genetic variation (e.g. Khila et al., 2009; Rowe et al., 2018; Crumière & Khila, 2019; Pruvôt et al., 2025).

## Endmatter

### Data and code availability

Data and code for this paper are available via a zenodo repository: https://zenodo.org/records/17520634? token=eyJhbGciOiJIUzUxMiIsImlhdCI6MTc2MjI1MTg3OSwiZXhwIjoxNzc1MDAxNTk5fQ.e yJpZCI6ImUyOTI1Y2E2LTNjMDctNDI5NC1hMjRlLTJlMTllYjc5OGEyZiIsImRhdGEiOnt9LC JyYW5kb20iOiI5YTA5NjhlYmNjNzlkYjFmMmQ0YjIzOTVhYWNjZjFiNyJ9.ZVOhIV3KmJh2h EwiQvRzEOCmx34oVDq6NOxwCr2OVvxJua1k4a2Px15ou3iPSblVvBa5jXAx50zVnqcMX Ukxxw

### Author contributions

R.A.W.W. contributed with the conception of the study, funding acquisition, data collection and analysis, and led the writing of the manuscript. A.T. contributed with experimental design, collection and analysis of data, R.R.S. contributed with the conception of the study, funding acquisition, data collection, and writing of the manuscript.

### Funding

This work was funded by a Starting Grant from the Swedish Research Council (Vetenskapsrådet) [grant no. 2022-03701], and a European Commission Marie Sklodowska-Curie Actions Independent Postdoctoral Fellowship [grant no. 101061664] to RAWW, as well as a Project Grant from the Swedish Research Council (Vetenskapsrådet) to RRS [2018-04598].

## Acknowledgements

We are grateful to the DrosEU and DrosRTEC consortia for their heroic efforts of sampling, sequencing, and establishing of isofemale line resources of *Drosophila melanogaster*, and also for helpful feedback. In particular we are grateful to Thomas Flatt, Pau Carazo, Mathieu Gautier for advice in the planning of experiments; to Élio Sucena for maintaining and shipping us isofemale lines. We are also grateful to Mirjam Amcoff, Nadeem Naber, Caja Samsen, and Matilde Stella for help in the lab during experiments and to Luc Bussière and Mike Ritchie for comments that improved the manuscript. The computations and data handling was enabled by resources provided by the National Academic Infrastructure for Supercomputing in Sweden (NAISS), partially funded by the Swedish Research Council through grant agreement no. 2022-06725.

