## Supplementary materials for "Sexually antagonistic co-evolution at the molecular level: patterns of genetic co-variation predict phenotypic outcomes of mating interactions"

**Thorson et al.**

### 1.Supplementary Methods

#### 1.1 Hierarchical Fst analysis

We obtain SNPs representing neutral or background genetic variation from short introns (length < 60 bp), which have previously been used to represent putatively neutrally evolving regions of the genome, and therefore a source of demographically informative SNPs (e.g.Kapun et al., 2020). We extracted intron coordinates from the *D. melanogaster* annotation (r6.12) and filtered them to include only autosomal introns < 60bp in length. Next, the full .vcf file of variants called by PoolSNP in the DEST v. 2.0 dataset was filtered to include only SNPs that occurred within these introns. This produced a final set of N = 44,838 SNPs in short introns. These SNPs were then used to conduct a hierarchical Fst analysis using poolfstat (Gautier *et al.*, 2013; Hivert *et al.*, 2018; Gautier *et al.*, 2024) using the locality of each sample as a grouping factor. Analysis showed significant among group Fst (Fgt = 0.04 [0.038, 0.045]), and within-group Fst which was low and close to 0 (Fsg = 0.01, [0.00962, 0.0104]). For each population, we therefore chose the sample with the highest average coverage for downstream analyses. We also exclude populations from the ancestral range in Africa (see the main text). The final set of population samples included N = 149 samples (Table S1).

#### 1.2. Definition of population-specific Fst

For each of the population genomic samples, we computed population-specific Fst measures for all SNPs within SFP loci and the set of randomly selected genomic background loci. Population-specific Fst was computed by a method following Weir & Goudet (2017) and Hivert et al.(2018) using the “poolfstat” R package (v. 2.1.2; Hivert *et al.*, 2018).

Fst is defined as:

$$Fst = (Q1 – Q2)/(1 – Q2)$$

For each population, Q2 values were obtained as the mean Q2 across all pairwise Q2 values from the compute.pairwiseFST() function of poolfstat (using method = “identity”). For each population, Q1 was computed as:

$$Q1_{pop} = 1 - (n\_hap/(n\_hap-1)) * ( ((r1*(r1-1))/((r1+r2)*((r1+r2)-1))) + ((r2*(r2-1))/((r1+r2)*((r1+r2)-1))) )$$

From the equation A37 in the supplementary materials of Hivert et al. (2018). Where *n_hap* is the number of haplotypes present in each pool, and *r1* and *r2* are the counts of the alternative and reference alleles at each SNP, respectively. One interpretation of this measure of Fst is thus the degree to which a population is different from the overall average, or in other words, the degree of exaggeration of allele frequencies compared to the overall average. This interpretation is useful in the analysis of molecular traits in the context of SAC. Note that Fst values can be negative if there are e.g. a large number of private alleles at high frequency.

#### 1.3. Computation of the relatedness matrix

We thinned all ~4.6 million SNPs in the DEST dataset to sets of 100,000 SNPs from across the genome. We used BayPass (v. 2.4; Gautier *et al.*, 2013) to run the core model and estimate the Ω matrix as well as the shape parameters of the beta distribution for $\pi_i$ (the ancestral allele frequency). We ran the core model for 11 independent sets of 100,000 SNPs and compared omega matrices and shape parameters across all runs to ensure that there was convergence. We checked that matrices were similar across runs using FMD distances, as in Gautier et al. (2013). Finally, we summarised the omega matrix by taking the mean value for each cell across runs.

#### 1.4. Isofemale line resources and stock population cages

We used isofemale lines from 6 natural European populations, available through the DrosEU consortium (Durmaz Mitchell *et al.*, 2025), and started 3 replicates for each natural population in population cages (dimensions 24.5 x 24.5 x 24.5 cm; BugDorm, MegaView Science Co., Ltd.). Each population cage was seeded with 5 males and 5 females from between 15 and 20 isofemale lines, depending on the natural population of origin (except for a single isofemale line of the MA population from which only 3 individuals for each cage could be obtained). Stock populations were allowed to mate freely and supplied with a bottle containing 65-70mL of fresh standard fly medium (in 1L $dH_20$: 80g cornmeal, 18g dried yeast, 10g soya flour, 80 malt extract, 40g molasses, 8g agar, 25mL 10% Nipagin, 4mL propionic acid) every 2 weeks. Populations were left to recombine for a minimum of 3-4 generations, and up to 9 generations, prior to the experiment. Cages were kept in incubators (Panasonic MIR-154-PE, Panasonic Healthcare Co. Ltd.; Termacks KB 8400L, Nordic Labtech AB) at 25º C and 12h:12h light:dark cycles. Cage placement within and between incubators was shuffled every generation at the time of food change.

#### 1.5. Statistical analyses

##### 1.5.1 Experiment 1

Re-mating rates were modelled as a function of the continuous measure of relative genetic differentiation between male and female components of the SP-network, where lower values indicate a female advantage, and higher values indicate a male advantage:

$$y \sim relative\ genetic\ differentiation + (1|population/replicate) + (1|batch) + e$$

Here we examined the hypothesis that there should be a negative relationship between the degree of relative genetic differentiation and re-mating rates across all populations. *e* is the residual error term modelled as Bernoulli trials with a logit link function.

We also fit models that used population of origin as a categorical predictor while controlling for replicate cages and experimental batch as random effects:

$$y \sim population + (1|population/replicate) + (1|batch) + e$$

To test for an effect of population, we fit a second model without the “population” term and compared model fits with the Leave One Out (LOO) approach. We also tested the specific contrast comparing the populations with inferred female advantage to those with inferred male advantage. We then tested whether the difference between female-advantage and male-advantage populations (see Results) was > 0:

*q1="((populationVA)+(populationMA))/2-((populationUM)+(populationGI))/2 > 0"*

Large positive effect estimates in this contrast thus indicate higher re-mating rates in VA and MA populations than in UM and GI populations.

Mortality/survival data was first modelled in a survival analysis using Cox proportional hazards model with population of origin, mating status, and their interaction as covariates, while controlling for replicate cages and experimental batch as random effects. Cox proportional hazards models were run using the “coxme” R package (v. 2.2-22; Therneau, 2024). We tested for differences between female and male advantage populations using specified contrasts with the “emmeans” package (v. 1.10.5; Lenth, 2025).

Data on reproductive success (the number of offspring) was modelled as a function of the continuous measure of relative genetic differentiation between male and female components of the SP-network, the mating status of the female (re-mated or not re-mated), and the interaction between mating status and the relative genetic differentiation. We also include the number of offspring in the first vial as a measure of the base-line fecundity of each female, although effects are qualitatively the same with or without this parameter (Figure S6).

y ~ *vial.1* + relative genetic differentiation + *mating_status* + *relative genetic differentiation:mating_status* + (1|population/replicate) + (1|batch) + e

We also fit models with population of origin as the predictor, along with mating status (mated once or re-mated), and their interaction, while controlling for replicate cages and experimental batch as random effects:

*y ~ vial.1 + population + mating_status + population:mating_status + (1|population/replicate) + (1|batch) + e*

*e* is the residual error term modelled as a normal distribution. First we modelled the response variable was either the number of offspring for each female (reproductive success). Because the number of offspring from vial 2 and vial 3 was strongly correlated with the total number of offspring produced across females (Figure S7), we here report only results from the offspring vials 2 and 3 in order to maintain a larger sample size (e.g. as females died as the experiment progressed). Results are qualitatively the same using the total number of offspring produced (Figure S8A). As above, we also include the number of offspring in the first vial as a measure of the base-line fecundity of each female, effects are qualitatively the same with or without this parameter (Figure S8B).

Next we modelled the log of the number of days until the vial with the maximum number of offspring (time to peak offspring production), in the same way as for offspring number. Here we remove females that produced no offspring at all (N = 16).

*y ~ vial.1 + relative genetic differentiation + mating_status + relative genetic differentiation:mating_status + (1|population/replicate) + (1|batch) + e*

As above, we also include the number of offspring in the first vial as a measure of the base-line fecundity of each female, but effects are qualitatively the same with or without this parameter (Figure S9). Not all females were included in these analyses due to loss from transferring into new vials after the re-mating assay.

We also modelled the time to peak offspring production as a function of population of origin, mating status (re-mated or not re-mated), and their interaction, while controlling for replicate cages and experimental batch as random effects:

*y ~ vial.1 + population + mating_status + population:mating_status + (1|population/replicate) + (1|batch) + e*

And, as above, we also tested whether the difference between female-advantage and male-advantage populations was > 0:

*q1="((populationVA)+(populationMA))/2-((populationUM)+(populationGI))/2 > 0"*

We always simplified models by LOO comparisons of nested models. With the exception of survival analyses, all models were implemented in a Bayesian framework using the “brms” R package (v. 2.22.0; Bürkner, 2017, 2018, 2021). With flat priors, MCMC chains were run for 10,000 iterations, after a warmup of 2,000 iterations with four independent chains. We checked chains for convergence visually.

**1.5.2 Experiment 2**

Survival was modelled using Cox proportional hazards model with population of origin as a fixed effect, while controlling for replicate cages and vial ID as random effects. Cox proportional hazards models were run using the “coxme” R package (v. 2.2-22; Therneau, 2024). We tested for differences between female and male advantage populations using specified contrasts with the “emmeans” package (v. 1.10.5; Lenth, 2025). We also analysed the data as lifespan, i.e. the time to death for each female. We modelled lifespan as function of population, including replicates and vials as random effects.

$$y \sim population + (1|population/replicate) + (1|population/replicate/vial) + e$$

We did not perform this analysis for survival data from experiment 1 because survival was overall very high, leading to very left-skewed data distributions.

**1.5.3 Statistical Software**

All statistical analyses were performed in R (v.4.4.2; R Core Team, 2025), in addition to the above named packages, several other packages were used in data-processing and analysis: “corrplot” (v. 0.95; Wei & Simko, 2024), “loo” (v. 2.8.0; Vehtari *et al.*, 2024), “coda” (v. 0.19.4-1; Plummer *et al.*, 2006), “tidybayes” (v. 3.0.7; Kay, 2024), “dplyr” (v. 1.1.4; Wickham *et al.*, 2023), “survminer” (v. 0.5.0; Kassambara *et al.*, 2025), “tidyverse” (v. 2.0.0; Wickham *et al.*, 2019), “lubridate” (v. 1.9.3; Grolemund & Wickham, 2011). All figures were plotted with “ggplot2” (v. 3.5.1; Wickham, 2016), “pammtools” (v. 0.5.93; Bender & Scheipl, 2018), “ggdist” (v. 3.3.2; Kay, 2025), and “cowplot” (v. 1.1.3; Wilke, 2024). We follow Muff et al. Muff *et al.*, 2022) and report results using the “language of evidence” for results of frequentist statistics and evaluate parameter estimates from Bayesian analyses in terms of the full posterior distributions and references to the 89% highest posterior density intervals (HPDIs; see McElreath, 2020).

### 3. Supplementary tables

See separate spreadsheet .xlsx file for supplementary tables.

### 4. Supplementary figures

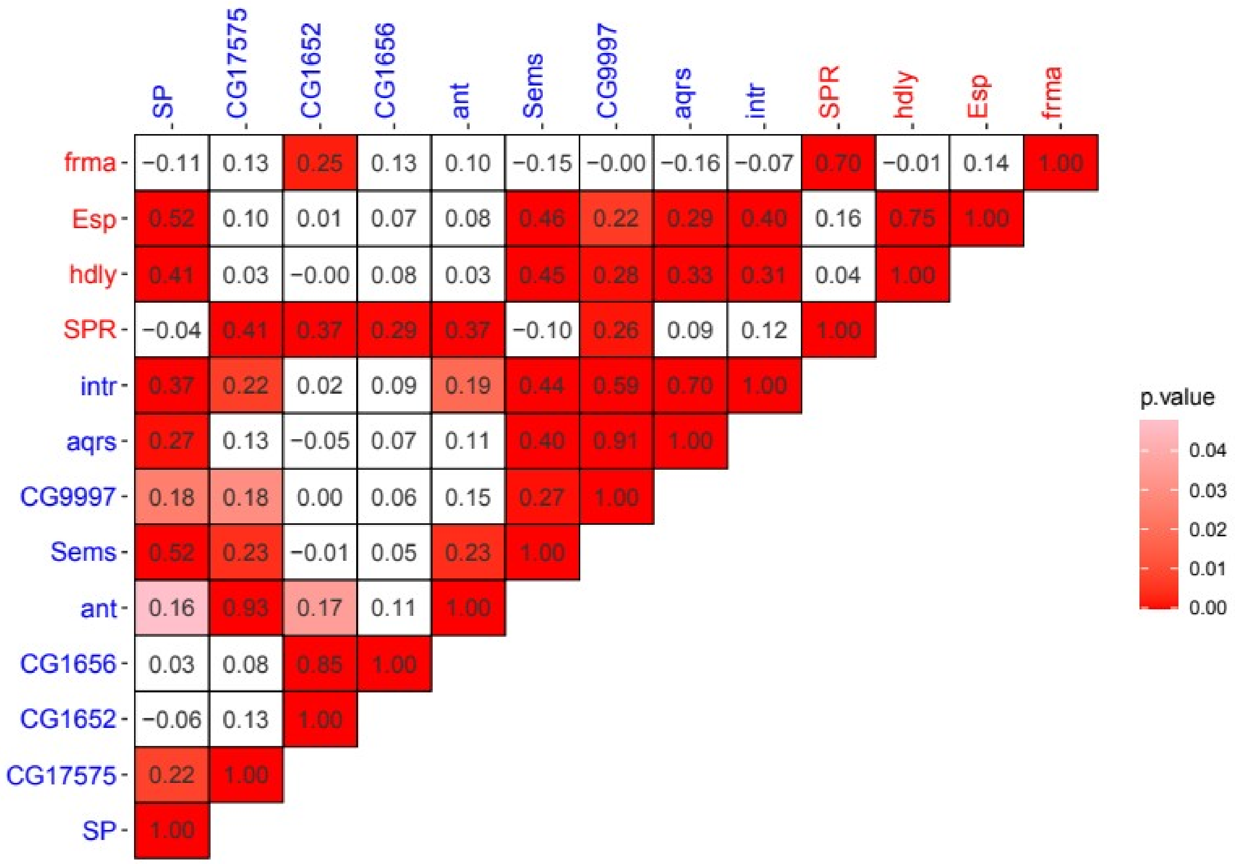

**Figure S1.** Pairwise Fst correlation matrix between all pairs of SP-network loci. Locus names are coloured red and blue for female- and male- components of the SP-network, respectively. The values in the cells give the Pearson’s correlation coefficient, and the colour of the cell reflects the p-value from a Pearson’s correlation test.

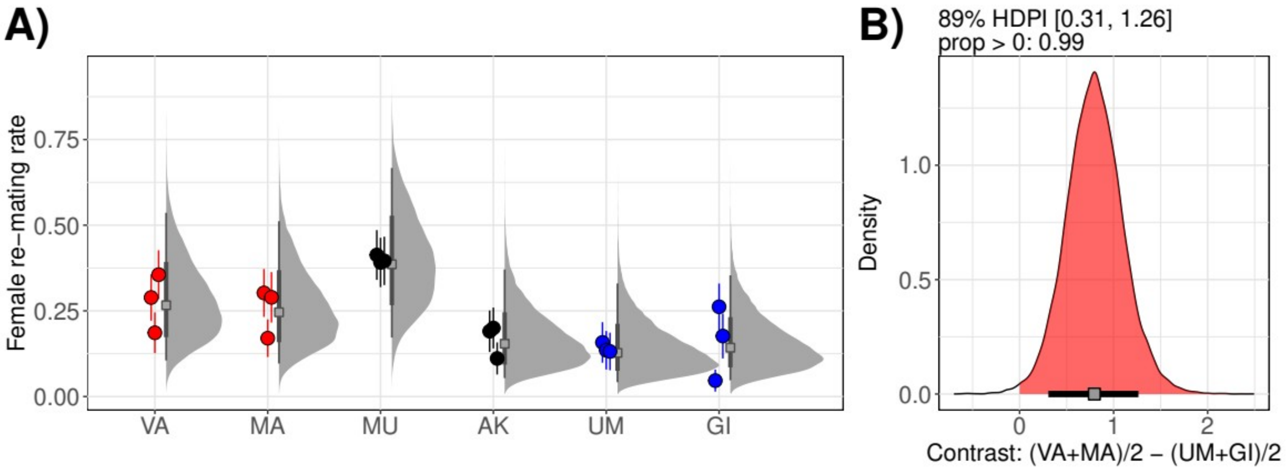

**Figure S2. A)** Female re-mating rates for each population. Empirical re-mating rates (±SE) are shown for each replicate of the six populations in points with error bars. Points are coloured by the sex with inferred advantage (red – female advantage - VA and MA; black – intermediate – MU and AK; blue – male advantage – UM and GI). Grey distributions, filled squares and intervals give the posterior distributions, mean, and 89% highest density posterior interval (HDPI) of predicted re-mating rates, respectively. **B)** The posterior distribution of predicted differences in re-mating rates between populations with inferred female advantage and inferred male advantage. The red filled area of the distribution gives the region > 0, the filled square and the interval line gives the mean and 89% HDPI of the difference. The numerical limits of the 89% HDPI and the proportion of the distribution > 0 are also given as text above the figure.

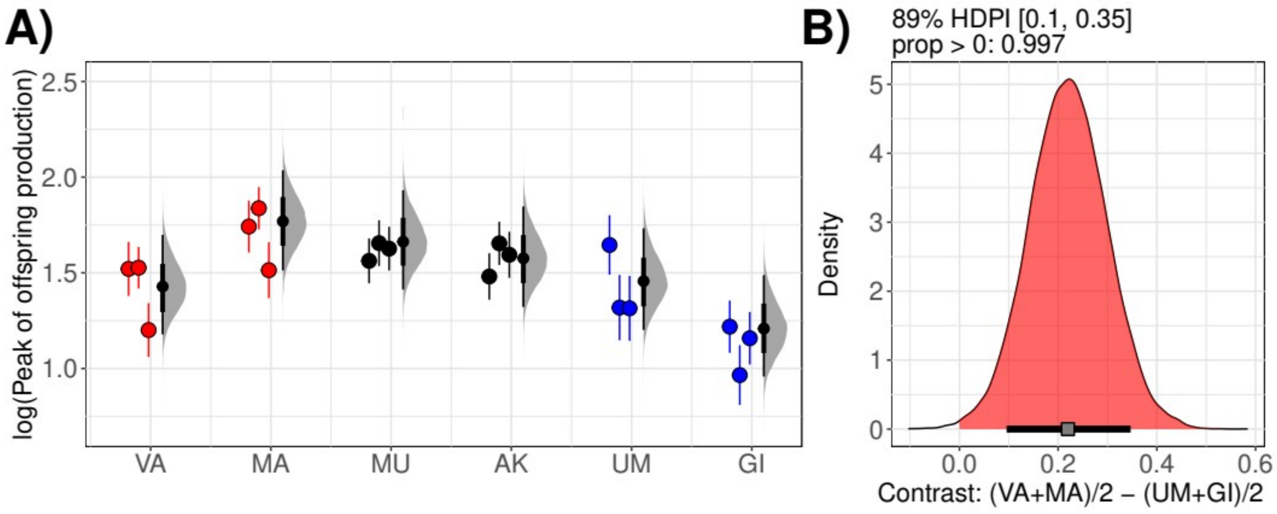

**Figure S3. A)** The log(number of days) until the peak of offspring production for each population. Empirical values (±SE) are shown for each replicate of the six populations in points with error bars. Points are coloured by the sex with inferred advantage (red – female advantage - VA and MA; black – intermediate – MU and AK; blue – male advantage – UM and GI). Grey distributions, filled squares and intervals give the posterior distributions, mean, and 89% highest density posterior interval (HDPI) of predicted re-mating rates, respectively. **B)** The posterior distribution of predicted differences in the The log(number of days) until the peak of offspring production between populations with inferred female advantage and inferred male advantage. The red filled area of the distribution gives the region > 0, the filled square and the interval line gives the mean and 89% HDPI of the difference. The numerical limits of the 89% HDPI and the proportion of the distribution > 0 are also given as text above the figure.

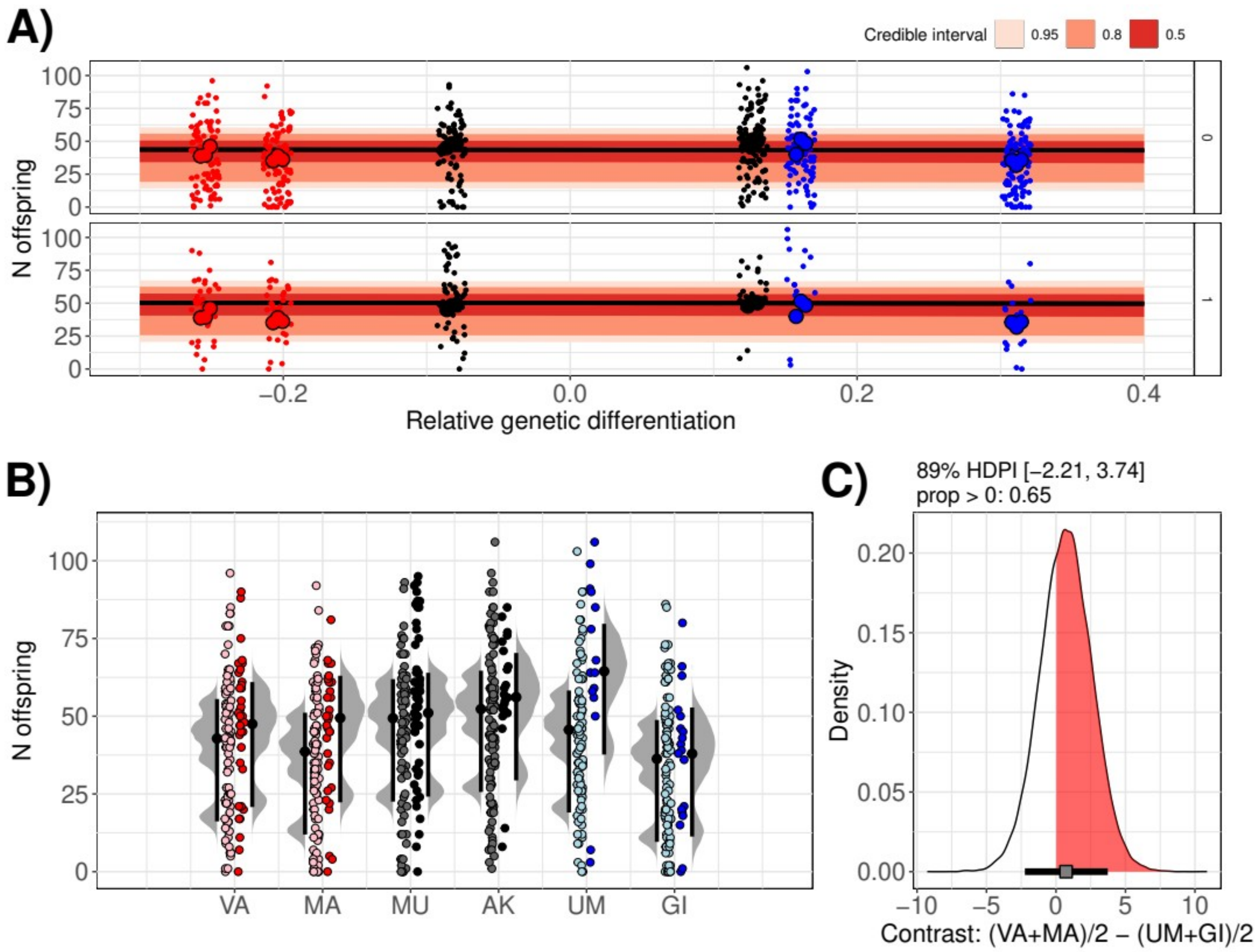

**Figure S4. A)** Points give the raw data for the number of offspring in vials 2 and 3 across populations. Data are split also by whether the female had re-mated (1), or not (0). Points are coloured by the sex with inferred advantage (red – female advantage - VA and MA; black – intermediate – MU and AK; blue – male advantage – UM and GI). The slope and error ribbons represent the posterior predicted mean and credible intervals for the slope. **B)** Points give the raw data for the number of offspring in vials 2 and 3 across populations. Data are split also by whether the female had re-mated (dark colours), or not (light colours). Grey distributions with points and error bars give the distribution of posterior predictions, along with the posterior mean prediction and the 89% HDPI. **C)** The posterior distribution of the contrast between female- and male-advantage populations. The red filled area of the distribution gives the region > 0, the filled square and the interval line gives the mean and 89% HDPI. The numerical limits of the 89% HDPI and the proportion of the distribution > 0 are also given as text above the figure.

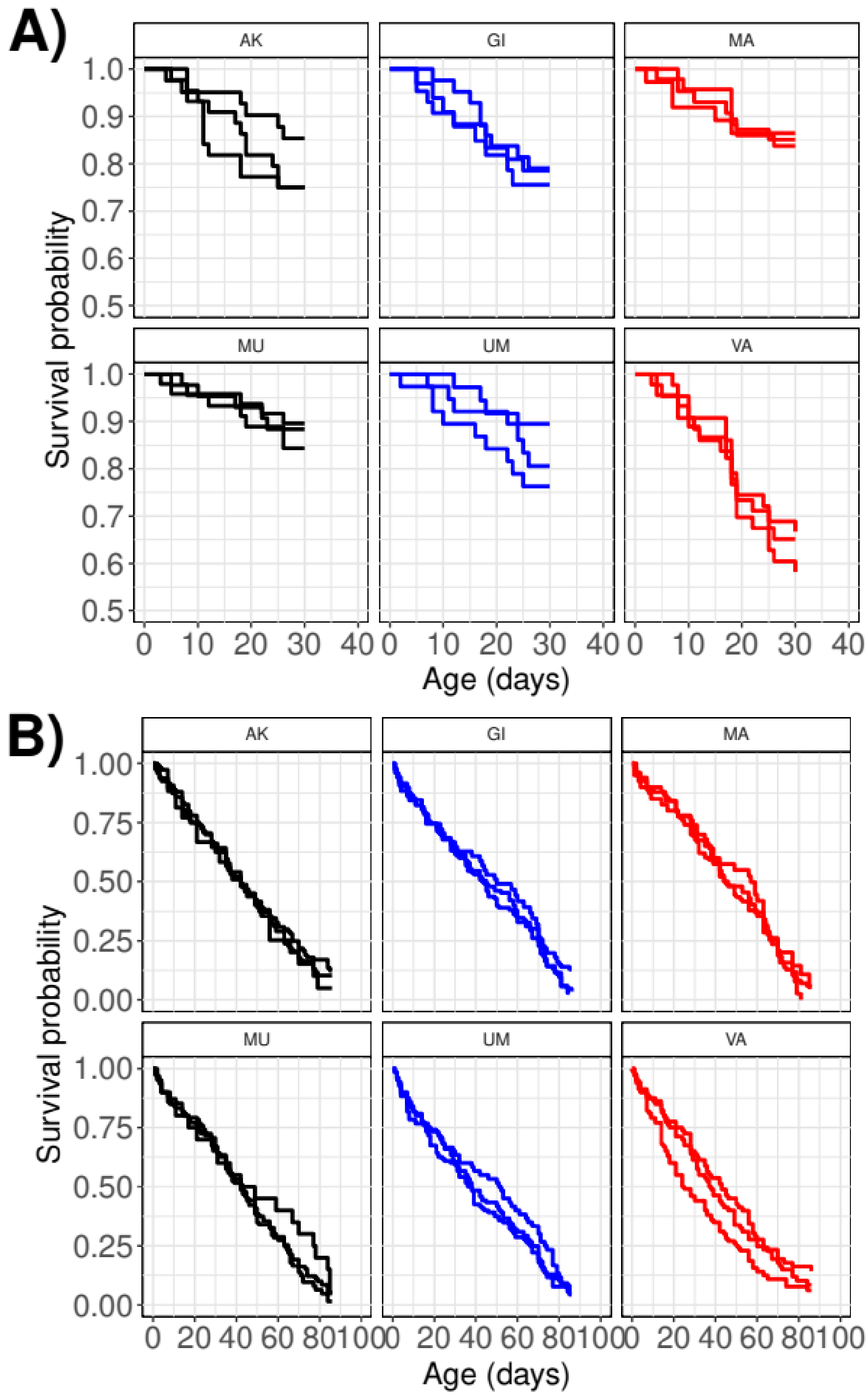

**Figure S5.** Survival curves for females from two experiments; **A)** tracking survival after a single re-mating opportunity and **B)** throughout life with continuous mating in groups of 5 females and 5 males. Lines are coloured according to which sex is inferred to have an advantage (red – female-advantage, blue – male advantage, black – intermediate).

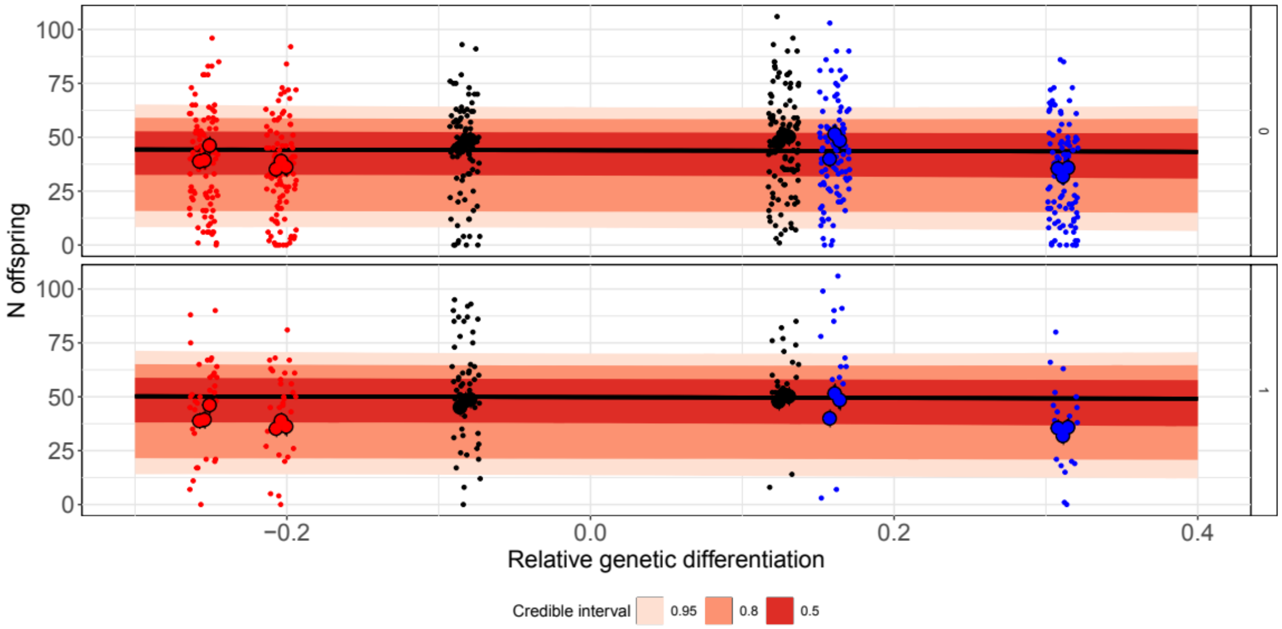

**Figure S6.** Points give the raw data for the number of offspring in vials 2 and 3 for each female across populations. Data are split also by whether the female had re-mated (1), or not (0). Points are coloured by the sex with inferred advantage (red – female advantage - VA and MA; black – intermediate – MU and AK; blue – male advantage – UM and GI). The slope and error ribbons represent the posterior predicted mean and credible intervals for the slope. Slopes are not different across panels ($ELPD_{w/\ interaction}$ -3244.0 ±20.9, $ELPD_{w/o\ interaction}$ -3243.0 ±20.9). The 89% HDPI for the effect of the relative genetic differentiation overlaps 0 (Estimate = -1.44, 89% HDPI [-14.74, 11.98]).

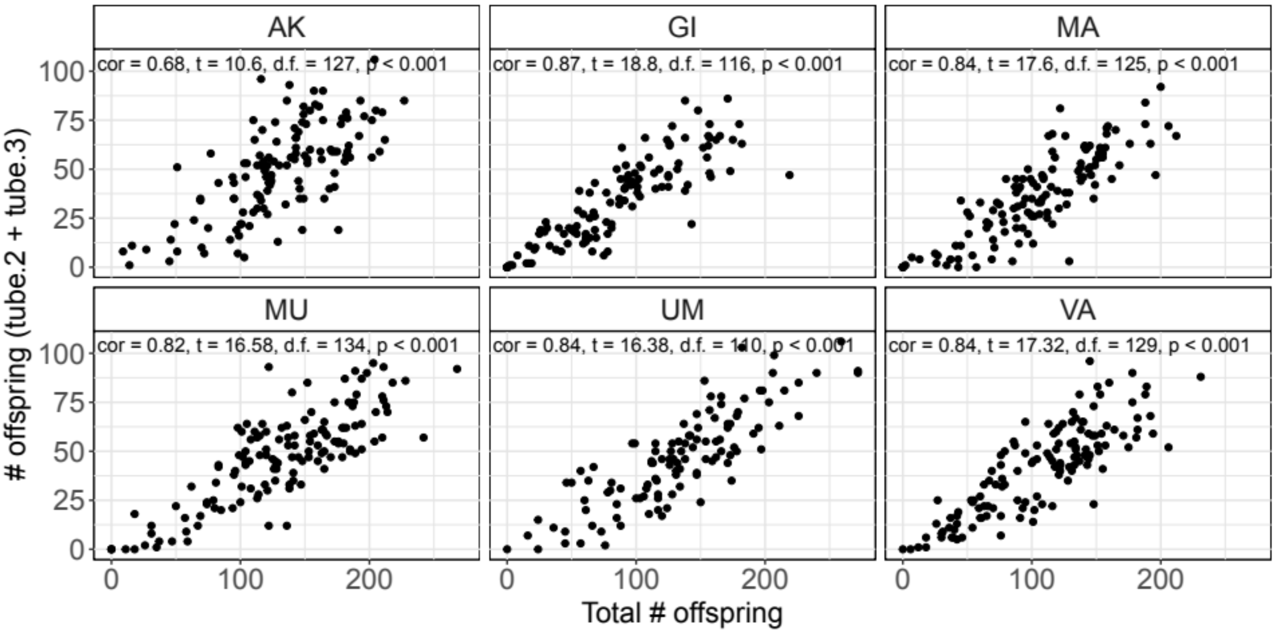

**Figure S7.** Correlation between the total number of offspring for each female, and the sum of offspring from vial 2 + vial 3. Inset text gives the results of a Pearson's correlation test for each population.

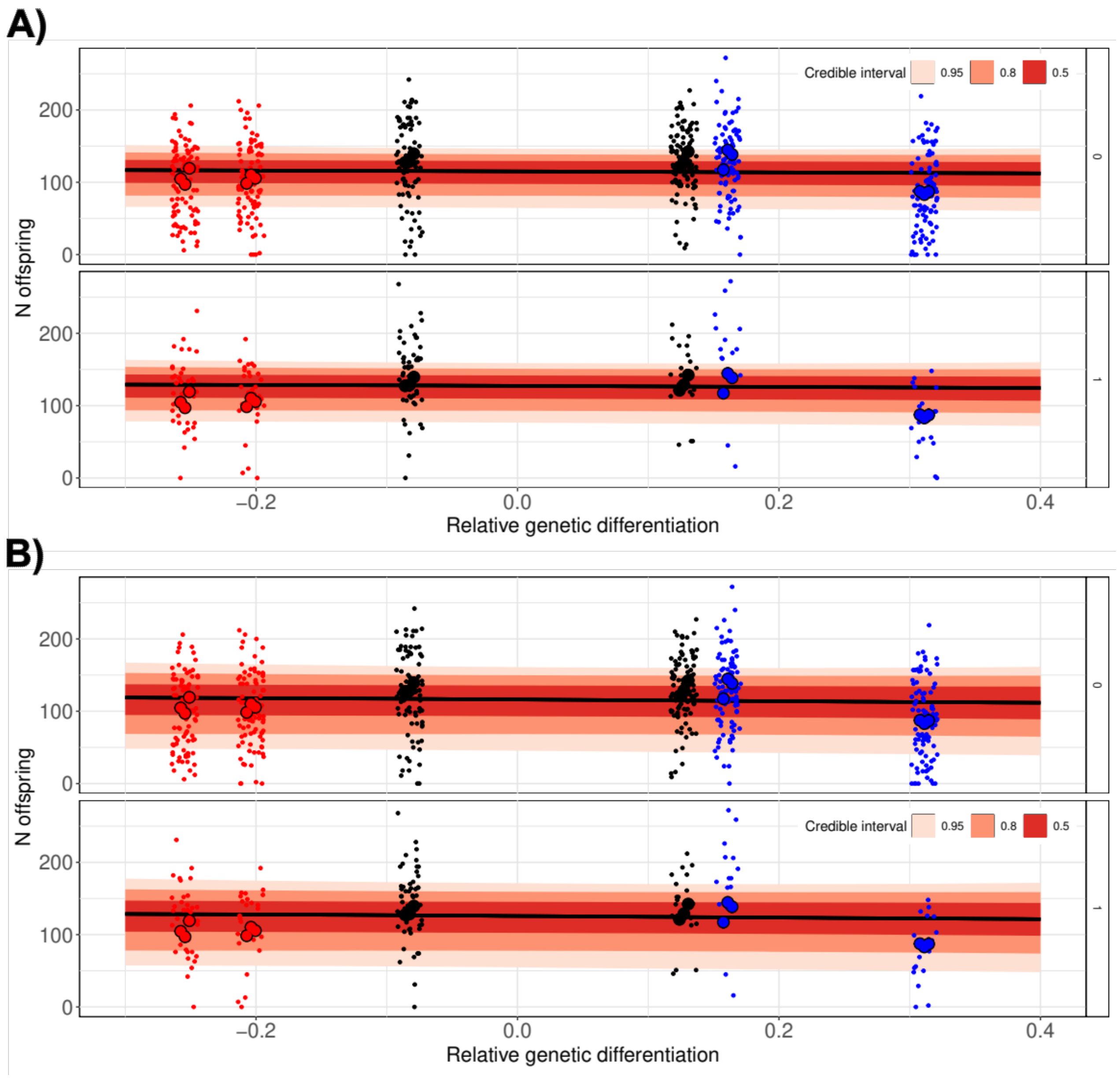

**Figure S8. A)** Points give the raw data for the total number of offspring for each female across populations. Data are split also by whether the female had re-mated (1), or not (0). Points are coloured by the sex with inferred advantage (red – female advantage - VA and MA; black – intermediate – MU and AK; blue – male advantage – UM and GI). The slope and error ribbons represent the posterior predicted mean and credible intervals for the slope. Slopes are not different across panels ($ELPD_{w/\ interaction}$ -3819.1 ±20.1, $ELPD_{w/o\ interaction}$ -3818.0 ±20.1). The 89% HDPI for the effect of the relative genetic differentiation overlaps 0 (Estimate = -5.73, 89% HDPI [-34.59, 23.02]). **B)** Same data as in **A)** but fitted model does not include the number of offspring in vial 1 as a co-factor. Slopes are not different across panels ($ELPD_{w/\ interaction}$ -3927.7 ±20.8, $ELPD_{w/o\ interaction}$ -3927.0 ±20.8). The 89% HDPI for the effect of the relative genetic differentiation overlaps 0 (Estimate = -7.76, 89% HDPI [-48.70, 32.71]).

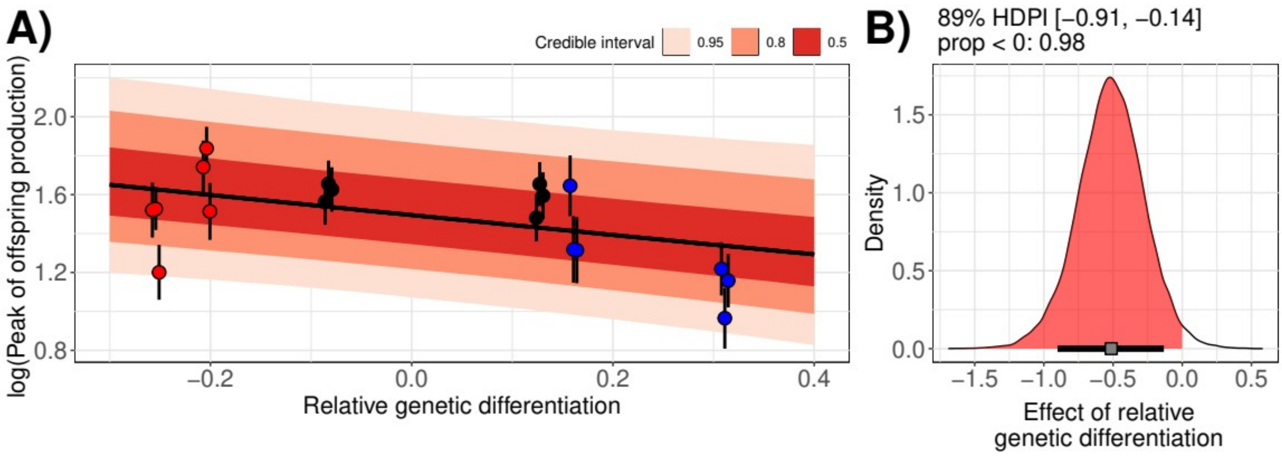

**Figure S9.** Same figure as in figure 3, but model does not include the number of offspring in vial 1 as a co-factor. **A)** The log(number of days) until the peak of offspring production as a function of a continuous measure of relative genetic differentiation at female- and male-components of the SP-network. Negative values indicate relatively more genetic differentiation in the female- *versus* the male-component. Each individual point is a replicate population cage, points are coloured by which sex is inferred to have an advantage in sexual conflicts (red – female-advantage, blue – male advantage, black - intermediate). The slope and error ribbons represent the posterior mean and credible intervals for the slope. **B)** Posterior distribution of the slope estimate. The area shaded in red indicates the proportion of the posterior distribution with slope estimates < 0, that supports a negative relationship between relative genetic differentiation and the peak of offspring production, the grey box and bar indicate the posterior mean and 89% HDPI respectively.
